# A GABA-A receptor response to CBD following *status epilepticus* in the medial entorhinal cortex of the rat

**DOI:** 10.1101/2024.10.23.619816

**Authors:** Benjamin S. Henley, Richard Walsh, Stuart Greenhill, Gavin Woodhall

**Affiliations:** Aston Institute of Health and Neurodevelopment, College of Life and Health Sciences, Aston University, Birmingham, UK; Department of Paediatric Neurosurgery, The Birmingham Women’s and Children’s Hospital NHS Foundation Trust, Birmingham, UK; University Hospital of North Durham, North Rd, Durham DH1 5TW

**Keywords:** cannabidiol (CBD), GABA receptor, NMDA receptor, interneurons, phytocannabinoids, temporal lobe epilepsy (TLE), anti-epileptic drugs (AEDs)

## Abstract

Cannabidiol is a non-psychoactive phytocannabinoid that has been implicated as a potential therapeutic in numerous neurological diseases. Perhaps the most widespread therapeutic use of CBD has been in the form of Epidiolex, which is used to treat seizures associated with Lennox-Gastaut syndrome, Dravet syndrome and tuberous sclerosis. Whilst the effectiveness of CBD in seizure reduction is clear, its mechanism of action is complex, and reflects the wide range of pharmacodynamic targets that includes receptors, ion channels and enzymes. This study investigated the effects of action of cannabidiol (CBD) on GABAergic transmission in layer II of the medial entorhinal cortex in animals that had previously undergone a period of *status epilepticus (SE)*. Spontaneous post-synaptic currents were recorded from medial entorhinal cortex layer II pyramidal cells in animals 1-7 after SE and in age matched controls as well as in tissue resected from children with temporal lobe epilepsy (TLE). CBD enhanced GABA_A_R-mediated inhibition by increasing decay times and inhibitory charge transfer across the postsynaptic membrane in status epilepticus (SE) but did not alter GABAergic transmission in age-matched control rats. The SE-induced effects of CBD were blocked by ligands acting as inverse agonists at the benzodiazepine site of the GABA_A_R receptor and the effects of CBD were additive to low-doses of benzodiazepine and barbiturate agonists, consistent with allosteric actions on the GABA_A_R. Similar effects were observed in both SE rat and human layer II neurons. Overall, these data suggest CBD may act as a positive allosteric modulator (PAM) at postsynaptic GABA_A_Rs and this action appears to develop following SE.

**Key points:**

- CBD increases inhibition only post *status epilepticus*.
- This effect is blocked by benzodiazepine site inverse agonists
- CBD is additive to low-dose zolpidem, suggesting an allosteric effect on the GABA_A_R
- CBD was similarly effective in *ex vivo* human epileptic tissue
- These results suggest CBD is a positive allosteric modulator at the GABA_A_R

## Introduction

Epilepsy has an incidence rate of roughly 1% of the global population (Leonardi and Ustun 2002) with Temporal Lobe Epilepsy (TLE) accounting for around 41% of all cases (Manford et al., 1992). Even in developed nations, many patients being treated for epilepsy suffer incomplete seizure control and/or resistance to commonly used anti-seizure medications (ASM; (Mao et al., 2024)). The prominent psychoactive component of cannabis, ^Δ9^-tetrahydrocannabinol (THC), is the most extensively studied phytocannabinoid. THC has shown some promise as an anticonvulsant, but its unwanted psychoactive effects preclude widespread medicinal use (Hill et al., 2012). On the other hand, the non-psychoactive phytocannabinoid, cannabidiol (CBD), has been shown to be a highly effective ASM (Cunha et al., 1980; Devinsky et al., 2016, Thiele et al., 2018) and in its formulation as Epidiolex, CBD is currently licensed for use in epilepsy in many nations including Europe and North America.

The pharmacology and mechanisms of action of CBD are complex. It is a low affinity allosteric antagonist at both subtypes of cannabinoid receptor (CB1R/CB2R; for review see Pertwee, 2008) and also acts as an antagonist at the GPR55 receptor (Pertwee et al., 2010). It has also been shown that CBD exerts is anticonvulsant effects through modulation of the effects of the endogenous membrane phospholipid, lysophosphatidylinositol (LPI), at GPR55 (Rosenberg et al., 2024).

Here we have investigated the anti-convulsant action of CBD in the RISE rat model of TLE (Modebadze et al, 2016) using spontaneous inhibitory postsynaptic currents (sIPSCs) as markers of inhibitory levels in layer II pyramidal cells in the medial entorhinal cortex (mEC). CBD enhanced inhibition in RISE but not in age-matched control rats. Similar responses also occurred in *ex vivo* epileptic paediatric human brain tissue. To explore how CBD reduced seizure activity and intensity we tested its effects on GABA_A_Rs using inverse agonists and benzodiazepine/barbiturate site agonists. Overall, our data suggest that anti-convulsant actions of CBD may be mediated by modulation of GABA_A_Rs.

### Animals and ethical approval

All procedures were carried out in accordance with the Animals Scientific Procedures Act (ASPA) 1986 UK, European Communities Council Directive 2010, and Aston University ethical review document. Human studies were conducted with tissue obtained with informed consent from pediatric patients undergoing epilepsy surgery at Birmingham Children’s Hospital. Ethical approval was obtained from the Black Country Local Ethics Committee (10/H1202/23; 30 April 2010), and from Aston University’s ethics committee (Project 308 cellular studies in epilepsy) and through the Research and Development Department at Birmingham Children’s Hospital (IRAS ID12287).

## Methods

### Brain slice preparation

Slices were prepared from male Wistar rats. Experimental objectives determined the age and strain of animal used. For local field potential (LFP) recordings, 450 μm thick slices were prepared. For whole cell patch-clamp recordings, 350 μm thick slices were prepared.

To prepare for brain extraction, each animal was first anaesthetised using 5% isoflurane in N2/O2. Once anaesthetised, the animals were injected subcutaneously (SC) with pentobarbital (600mg/kg), and intramuscularly (IM) ketamine (100mg/kg) and xylazine (10mg/kg) to induce terminal anaesthesia and neuroleptanalgesia. Correct depth of anaesthesia was observed by absence of normal paw pinch and corneal reflex. Next, the animals were bathed in ice-cold water for ∼45 seconds before the transcranial perfusion was performed using 20 – 40ml ice-cold sucrose-based artificial cerebrospinal fluid (aCSF, cutting solution) containing (in mM): 180 sucrose, 2.5 KCl, 10 MgSO4, 1.25 NaH2PO4, 25 NaHCO3, 10 glucose, 0.5 CaCl2, 1 L-ascorbic acid, 2 N-acetyl-L-cysteine, 1 taurine and 20 ethyl pyruvate. Neuroprotectants were also added to the cutting solution to improve slice viability, including (in mM): 0.045 indomethacin, 0.2 aminoguanidine, 0.4 uric acid, 0.13 ketamine and 0.2 brilliant blue G. The complete cutting solution was saturated prior to perfusion with 95% O2/5% CO_2_ (carbogen), pH 7.4, 300 – 310 Osm/L.

Follow rapid dissection and extraction of the brain, the brain was placed in the remaining cutting solution for transportation. Slicing was performed at room temperature in the ice-cold cutting solution using a 7000smz-2 model Vibrotome (Campden Instruments Ltd). Horizontal or coronal slices were made to contain the hippocampus/mEC or mPFC, respectively. Slices were then transferred to the holding chamber and stored at room temperature for 1 hour in a standard aCSF containing (in mM): 126 NaCl, 2.5 KCl, 1 MgCl2, 2.5 CaCl2, 26 NaHCO3, 2 NaH2PO4 and 10 D-glucose. Neuroprotectants were also added (in mM): 0.045 indomethacin and 0.4 uric acid. Continual perfusion of 95% O2/5% CO_2_ (carbogen) helped maintain pH 7.3, 300 – 310 Osm/L.

### *In vitro* electrophysiology

Whole cell patch-clamp experiments were conducted to understand inhibitory activity in the frontal and temporal lobes associated with epileptogenesis. Briefly, borosilicate capillary glass micropipettes were pulled using a P-1000 Micropipette Puller (Sutter Instruments, USA) with a tip resistance of 3 – 6 MΩ. Depending on the objectives of the experiments, these electrodes were filled with an internal solution specific for recording spontaneous inhibitory post-synaptic currents (sIPSCs). Composition of sIPSC solution (in nM): 100 CsCl, 40 HEPES, 1 QX-314, 0.6 EGTA, 5 MgCl_2_, 10 TEA-Cl, 80 ATP-Na, 6 GTP-Na, 1 IEM1460 final osmolality of 285 mOsmol and pH 7.3.

Following a one-hour acclimatisation period, slices were transferred to the patch clamp rig and recordings were performed in a submerged chamber. Temperature of the circulating ACSF was maintained at ∼30 °C using a TC-324C Single Channel Temperature Controller (Warner Instruments, USA). Filled electrodes were attached to a CV 203BU head stage (Molecular Devices, USA) which was controlled using PatchStar Micromanipulator (Scientifica, UK). Basler Ace 2.3 MP PowerPack Microscopy camera (Basler, Germany) is used for visualisation of target cells. Once an appropriate cell was identified and the electrode is lowered to just above the cell, a seal was formed between the membrane and the electrode, and a resistance of at least 1 GΩ (GigaOhm) was achieved before breaking through. Upon breaking through, a seal test (5 mV pulse) was applied for 10 minutes to allow for adequate filling of the cell with the internal solution and space-clamp. An Axopatch 200B Microelectrode Amplifier (Molecular Devices, USA) was used to hold cells at -70 mV. After filling, at least 15 minutes of baseline was recorded before the bath application of drugs. A seal test was applied every 5 minutes to monitor any changes in the access resistance. Any recordings where the seal test altered >30% of the original access resistance were discarded.

Data were acquired using Clampex 10.7 software with a sampling rate of 10 KHz, and filtered at 2 KHz which was digitised using a Digidata 1440A. Data was analysed using Clampfit 10.7, GraphPad prism 8 and Microsoft Excel.

### Reduced Intensity Status Epilepticus model of epilepsy and epileptogenesis

The Reduced Intensity Status Epilepticus (RISE) model of epilepsy and epileptogenesis was created in our laboratory (Modebadze et al., 2016) as a low mortality, high morbidity modification of the pilocarpine model. In short, 24 hours prior to induction, the animals were treated with lithium chloride (LiCl, 127 mg/kg, SC). On the day of induction, α-methyl scopolamine (1mg/kg, SC) was administered to reduce the peripheral effects of muscarinic cholinergic receptor activation with pilocarpine. Thirty minutes later, low dose pilocarpine (25 mg/kg, SC) was administered. Animals were then closely monitored for seizure activity using the Racine’s scale (Racine, 1972) as previously described (Modebadze et al., 2016; Needs et al., 2019).

Those rats who progress to stage 5, recovered and re-entered stage 5 a second time were considered to have entered status epilepticus (SE). Animals then remained in modified SE for no more than one hour, at which point seizure activity was terminated using standard methods (Modebadze et al., 2016). Behavioural signs of SE ceased within 30 minutes and animals were closely monitored while allowed to recover on a heat pad with treats. 1 ml 0.9% saline with glucose SC also given to rehydrate animals. In most cases, animals regained normal behaviour within a few hours, with complete recovery within 12 hours. Brains were taken for *in vitro* investigation between days 1-7 post *status*.

### Human tissue experiments

Specimens were resected intraoperatively with minimal traumatic tissue damage, and minimal use of electrocautery. For transport to the laboratory, samples were transferred immediately to ice-cold choline-based artificial cerebrospinal fluid (aCSF) standardized for use in human tissue experiments containing in mmol/L: 110 choline chloride, 26 NaHCO_3_, 10 D-glucose, 11.6 ascorbic acid, 7 MgCl_2_, 3.1 sodium pyruvate, 2.5 KCl, 1.25 NaH_2_PO_4_, and 0.5 CaCl_2_ with added 0.04 indomethacin and 0.3 uric acid for neuroprotection, and bubbled with carbogen (95% O_2_, 5% CO_2_). For slice storage and experiments, aCSF containing (in mmol/L): 125 NaCl, 3 KCl, 1.6 MgSO_4_, 1.25 NaH_2_PO_4_, 26 NaHCO_3_, 2 CaCl_2_, 10 Glucose, was used.

### Statistical analysis

200 sIPSCs were sampled for each condition in each recording, and median values were calculated prior to comparison using the Mann-Whitney test (2-tailed, alpha = 0.05) to compare mean medians.

### Source of drugs used in this study

Drugs used in this study were supplied by Sigma-Aldrich, UK.

## Results

### CBD effects on SE and AMC rat mEC layer II pyramidal cell sIPSCs

We first investigated the effects of CBD on status epilepticus (SE; n = 26; animal n = 19) and age-matched control (AMC; n = 12; animal n = 9) rats using voltage-clamp of mEC Layer II pyramidal cells (raw traces shown in **Figure 1A**). Quantitative analyses of these data (**Figure 1B**) showed that CBD did not change the mean median amplitude in control rats. **Ai** and **Bi** show the mean median amplitude for each condition in the control sIPSC vs 30μM CBD. (**Ai**; AMC; 29.9 ± 9.3 vs 34.8 ± 11.5; p = 0.11). (**Bi**; SE; 26.8 ± 2.3 vs 29.8 ± 3.1; p = 0.12). **Aii** and **Bii** show that CBD did not change the mean median IEI (ms) in control rats (**Aii**; AMC; 112 ± 17.8 vs 120 ± 25.1; p = 0.54). In contrast, a significant decrease in mean median IEI was observed in epileptic rats (**Bii**; SE; 77.8 ± 6.6 vs 71.9 ± 5.7; p = 0.02). There was no change in the mean median decay time constant (*tau*, ms) following CBD application in the control rats (**Aiii**; AMC; 8.99 ± 0.97 vs 9.20 ± 1.06; p = 0.39;). There was, however, a significant increase in the epileptic rats (**Biii**; SE; 8.62 ± 0.5 vs 9.30 ± 0.68; p = 0.045). Finally, the normalised inhibitory charge transfer (NICT) values were unchanged in control rats by CBD (**Aiv**; AMC; mean value of 115 ± 12.4; range: 50.3-179; p = 0.27). In the SE rats a significant increase in NICT was noted following CBD application (**Biv**; SE; 131 ± 11.7; range: 31.7-727; p = 0.012).

**Figure 1.**
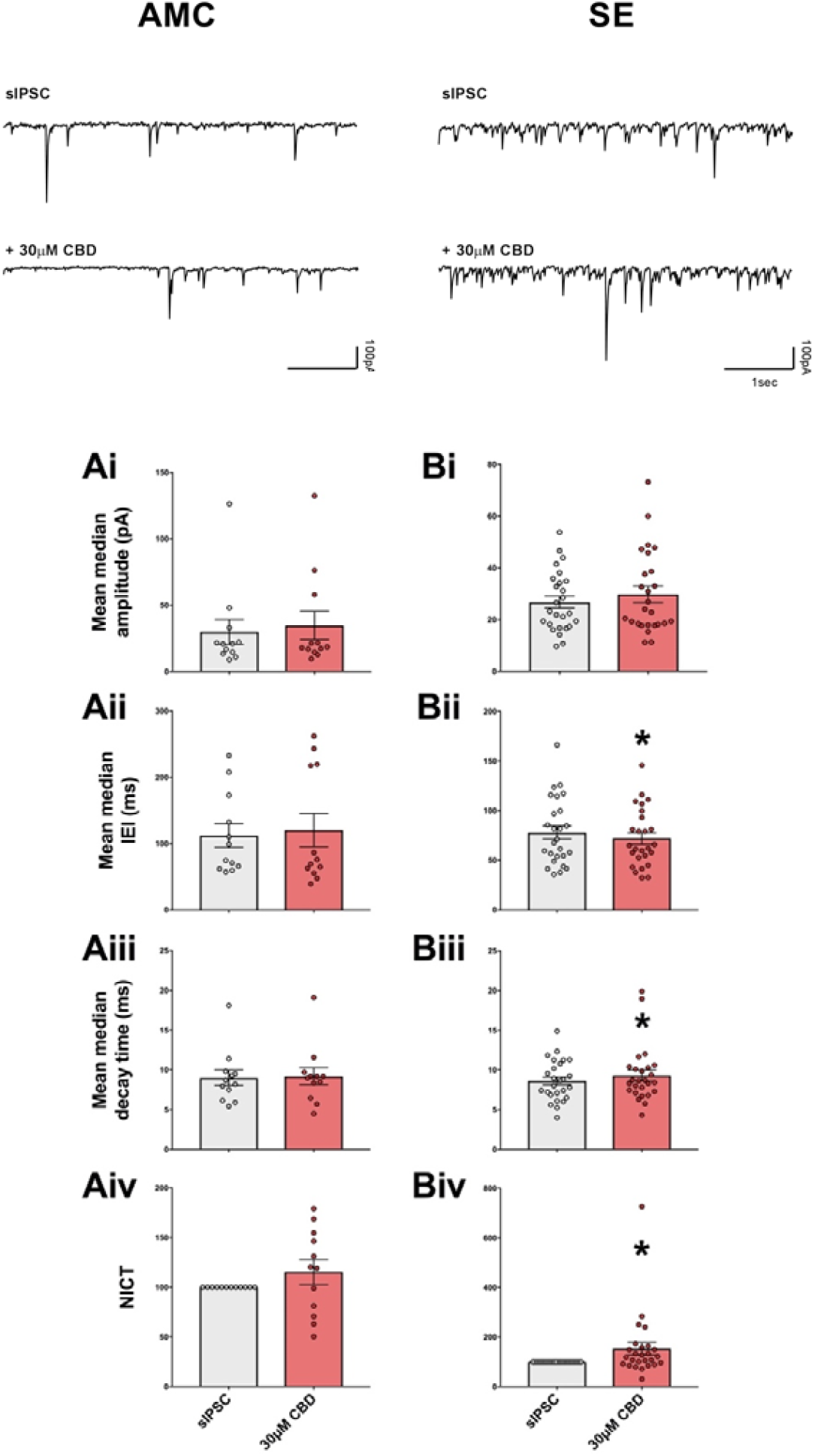
Effect of CBD on AMC and SE rats. **A**. sIPSC raw traces before and after 30μM CBD addition. Representative raw traces taken from age-matched controls (AMC; left; n = 12; animal n = 9) and status epilepticus (SE; right) rats, before and after CBD addition. **B**. Figure 2. Histograms showing quantification the effect of CBD on each of the sIPSC parameters. Ai/Bi show mean median amplitude for each condition in the control sIPSC and 30μM CBD. Aii/Bii represent mean median IEI comparisons (ms). Aiii/Biii highlight mean median decay time comparisons (ms). Aiv/Biv show comparisons of NICT values. AMC left, SE, right. * p < 0.05. See text for details.

We also tested the effects of 30μM CBD on pyramidal cell sIPSCs in human tissue slices obtained from paediatric patients undergoing surgery for refractory epilepsy. Representative traces are shown in (**Figure 2A**) with quantitative analyses in (**Figure 2B**; n = 9; sample number n = 7). Neither sIPSC amplitude nor the IEI changed on addition of CBD. However, consistent with rodent data, CBD did elicit a significant increase in decay time (ms) (**Aiii**; 8.08 ± 0.67 vs 9.31 ± 0.95; p = 0.009) and NICT values (**Aiv**; 148 ± 25 %; p = 0.039).

**Figure 2.**
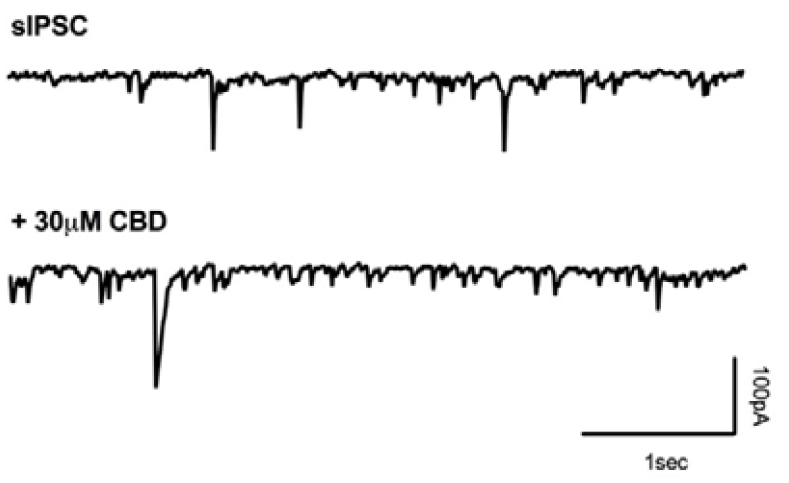
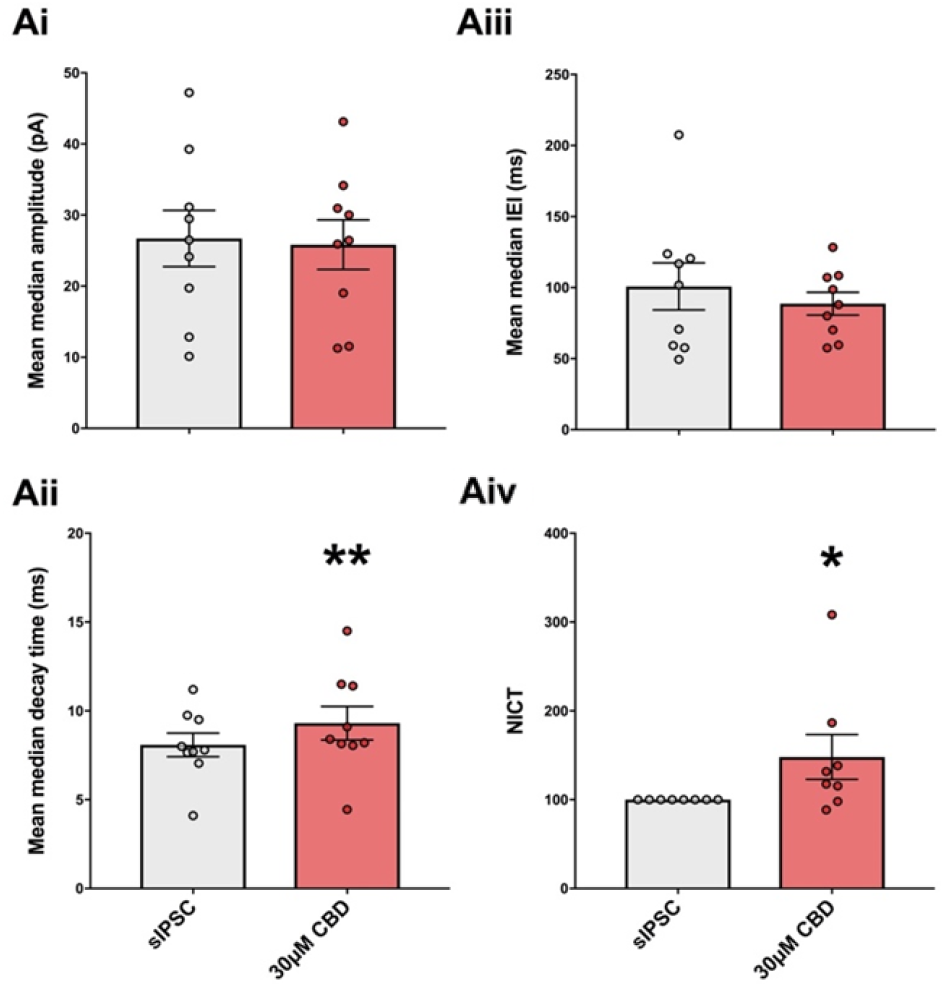
Effects of CBD in human tissue. A, Representative sIPSC rtraces from human pyramidal cell sIPSCs in response to CBD. Raw traces before and after 30μM CBD addition in a HT pyramidal cell. B, Quantitative analyses of responses of human neurons to CBD. Mean median amplitude (Ai), IEI (Aii) and decay time comparisons (Aiii), as well as NICT analysis (Aiv) for all samples analysed. * p < 0.05, ** p < 0.01.

Changes in the decay time (approximately 15%), coupled with lack of clear change in IPSC amplitude are consistent with actions of low doses of zolpidem at the benzodiazepine site on the GABA-A receptor elicited by zolpidem (for example see DeFazio and Hablitz, 1998). Since only slices from SE rats showed any CBD effect, GABA_A_R sIPSCs from SE rats were measured in the presence and absence of 30*µ*M CBD following 15 min prior application of the BZ inverse agonists 500nM flumazenil (n = 7, animal n = 5) and, separately, 5*µ*M β-carboline-3-carboxylic acid N-methylamide (β-carboline) (n = 5, animal n = 3). Representative raw traces of which are shown in **Figure 3A**.

**Figure 3.**
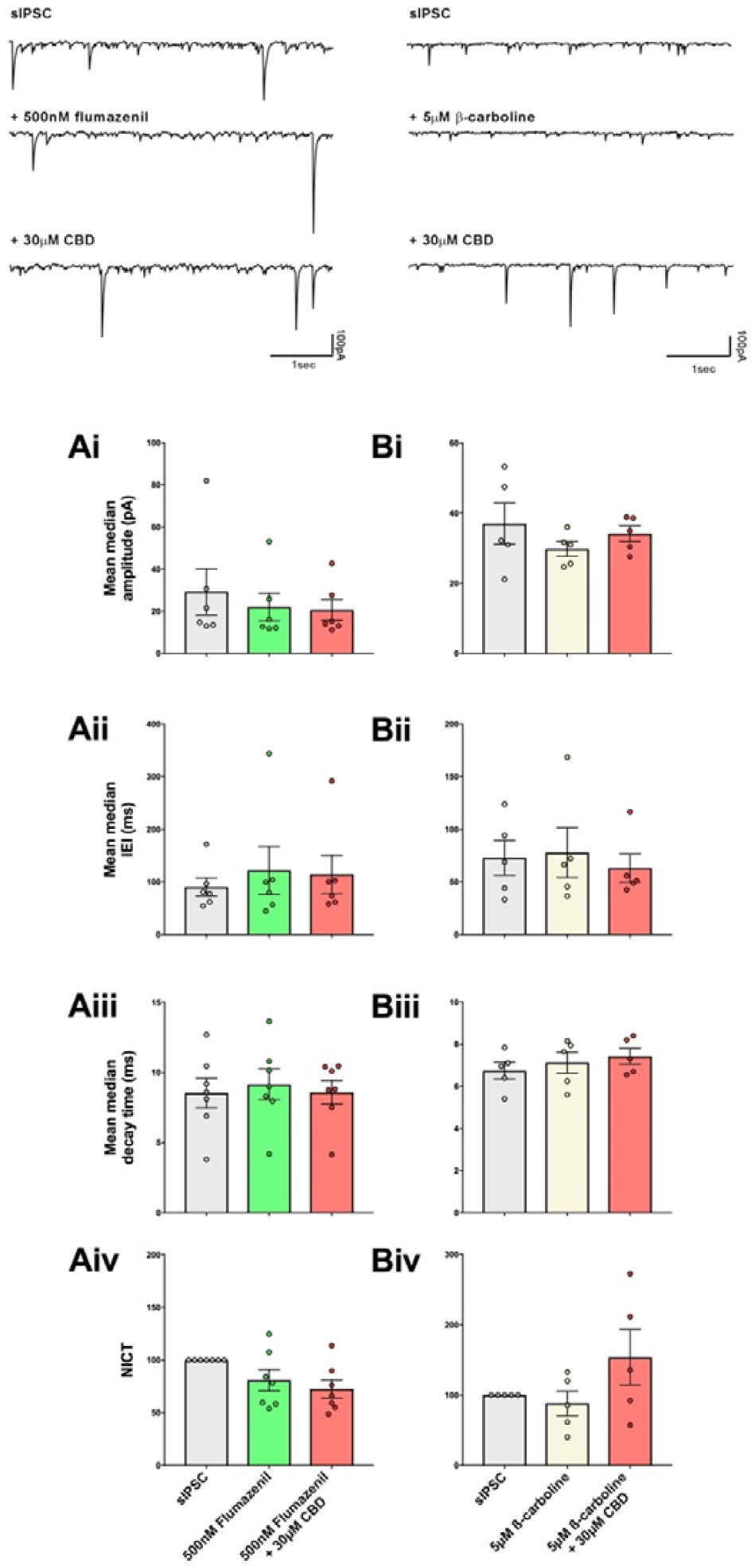
A. Raw sIPSC traces from SE rat tissue. Left 500nM flumazenil, right 1*µ*M β-carboline. B. Quantitative analyses of sIPSCs following application of the BZ site inverse agonists flumazenil (Ai-iv) and β-carboline (Bi-iv) prior to CBD. Mean median amplitude (Ai/Bi), IEI (Aii/Bii) and decay time (Aiii/Biii) and NICT values (Aiv/Biv). See text for details.

Analyses of sIPSCs ± flumazenil ± CBD are shown in **Figure 3B** (**Ai-iv**). Consistent with previous data from our lab (Prokic et al., 2015). Flumazenil had no effect on amplitude, IEI, decay times or NICT values. Mean median amplitudes were 28.8 ± 9.2 pA in control conditions and 22.4 ± 5.6 pA in FLZ and 20.5 ± 4.2 pA in FLZ+CBD; p > 0.05; **Ai**). However, the mean median IEI responses in slices from these epileptic rats were unchanged by flumazenil or flumazenil + CBD additions (**Aii**; control 86.0 ± 15.3 vs 115 ± 39.0 in FLZ and 110 ± 31.1 in CBD; p > 0.05). The mean median decay time constant did not increase on addition of flumazenil or CBD (**Aiii**; control 8.5 ± 1.0 vs FLZ 9.1 ± 0.8 and 8.6 ± 0.9 in FLX + CBD; p = 0.051). There were no changes in the mean NICT values for any drug condition (**Aiv**; control 80.9 ± 10.2 fC; CBD 72.6 ± 8.6 fC > 0.05). In order to confirm the results with FLZ, we performed experiments using the β-carboline alone and with addition of CBD. Experiments revealed a similar blockade of CBD effects in the presence of β-carboline (**Bi**; control amplitude 37.0 ± 5.8 pA vs β-carboline 29.8 ± 2.1 pA vs 34.1 ± 2.2 pA in β-carboline + CBD; p > 0.05). Similarly, no change in mean median IEI was noted (**Bii**; control 72.9 ± 16.5 vs 77.9 ± 23.6 in β-carboline and 63.1 ± 13.5 in β-carboline + CBD p > 0.05) or in mean median decay time constant (**Biii**; β-carboline 6.74 ± 0.41 – 7.12 ± 0.50; range: 5.40-7.85; 5.60-8.15; β-carboline + CBD; 7.43 ± 0.38; range: 6.55-8.40; p > 0.05). Similarly, the relative NICT values were unchanged by β-carboline (87.9 ± 17.4; range: 39.9-133). Although there appeared to be an increase in NICT values in the β-carboline + CBD conditions this was not statistically significant due to the large variance of responses (154 ± 39.4; range: 56.9-273; p > 0.52), with no significance shown in the post-hoc analysis.

We next tested the effects of the BZ site agonist, zolpidem, in SE rat neurons. We used both a low 100nM dose (n = 6, animal n = 5) and a higher 1*µ*M dose (n = 6, animal n = 5) of zolpidem ± CBD to investigate if they might have additive or occlusive effects respectively. Zolpidem (100nM) ± CBD had no significant effect on sIPSC amplitude or IEI responses (**Figure 4**). 100nM zolpidem alone, however, did increase sIPSC decay time whereas 100nM zolpidem + CBD did not show a significant difference from untreated. Similarly, 1*µ*M zolpidem did not significantly alter the sIPSC amplitude, IEI or decay times, but, as with 100nM, 1*µ*M zolpidem alone evoked a highly significant increase in the mean NICT value which decreased back to baseline levels on addition of CBD.

**Figure 4.**
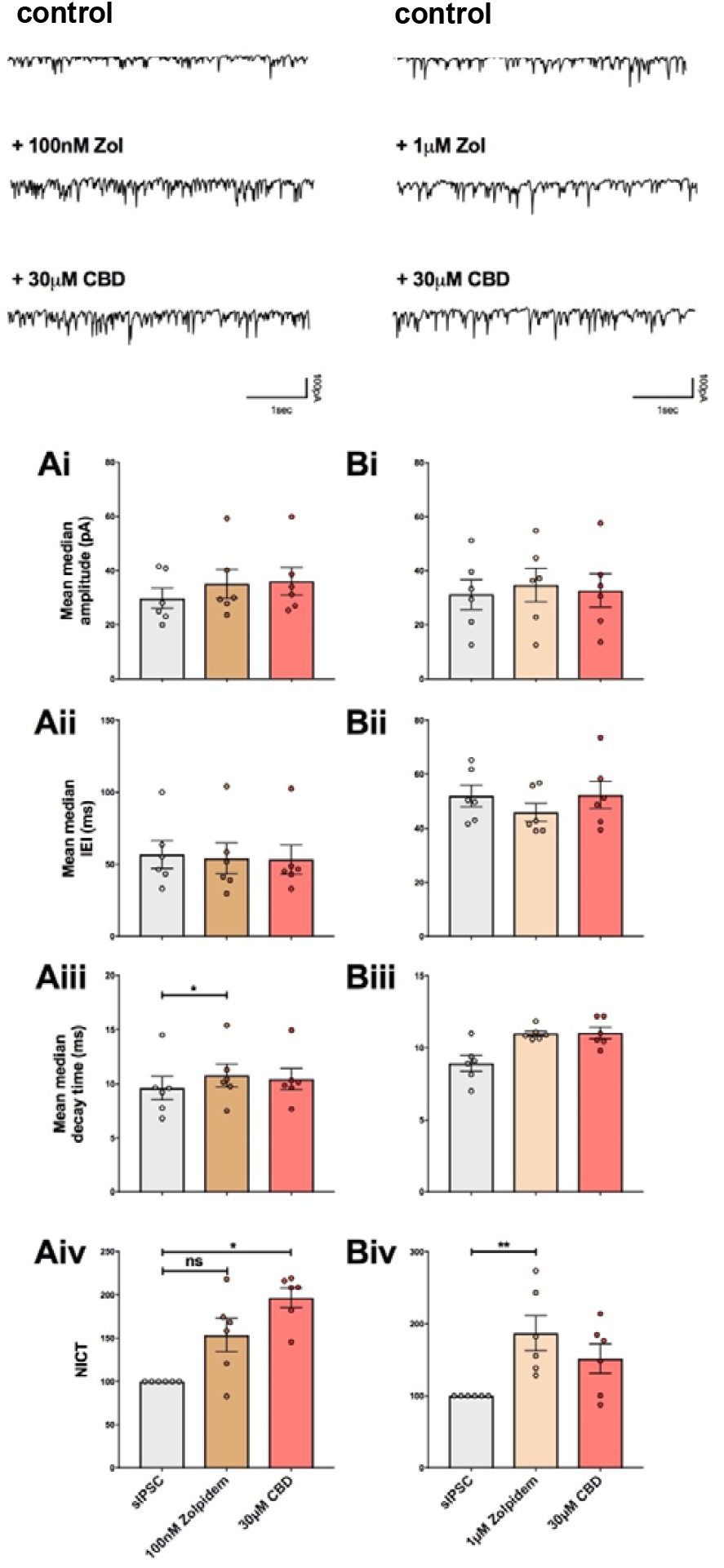
A. Raw traces of sIPSC from 100nM, 1*µ*M zolpidem and 40*µ*M pentobarbital treatments. B. Histograms for BZ/barbiturate site agonists effect on SE tissue. Parameters analysed for the addition of BZ and barbiturate agonists, being 100nM zolpidem (A), 1μM zolpidem (B) and, 40μM pentobarbital (C) including Ai-Ci: mean median amplitude; Aii-Cii: mean median IEI; Aiii-Ciii: mean median decay time; Aiv-Civ: NICT values. * p < 0.05, ** p < 0.01.

In neurons from SE slices, 100nM zolpidem had no significant effect on the mean median amplitude of sIPSCs (**Ai**; control 29.8 ± 3.8 vs ZOL 35.0 ± 5.3 vs 36.0 ± 5.2 in ZOL+CBD; p > 0.05) or the mean median IEI responses (**Aii**; control 57.0 ± 9.6 vs ZOL 54.1 ± 10.8 vs 53.2 ± 10.1 ms I ZOL+CBD; p > 0.05). 100nM zolpidem alone, however, did increase the mean median decay time constant (**Aiii)** from 9.6 ± 1.1 to 10.8 ± 1.1 ms). Surprisingly, 100nM zolpidem + CBD did not show a significant difference from untreated (**Aiii**; 10.4 ± 0.99; range: Relative NICT values are presented in **Aiv**, with a significant increase observed in only the CBD condition (100nM zolpidem: 154 ± 19.1; range: 82.8-218; CBD: 197 ± 11.5; range: 146-219). A p value of 0.012 was returned overall, with significance observed only in baseline vs. 100nM zolpidem + 30μM CBD (p = 0.012).

The initial zolpidem experiments at 100 nM suggested occlusion of the CBD effect by low-dose ZOL, however an overall increase in NICT was clear in the comparison between control values and the ZOL+CBD combination. We sought to fully occlude the CBD effect through use of a higher concentration of ZOL. At 1*µ*M zolpidem neither the mean median amplitude responses from SE rats (**Bi**; zolpidem - 31.2 ± 5.56 – 34.8 ± 6.18; range: 12.6-51.2; 12.6-54.8; p > 0.05) (zolpidem + CBD; 32.7 ± 6.18; range: 13.7-57.6; p > 0.05), nor the mean median IEI responses (**Bii**; zolpidem - 52.0 ± 3.95 – 45.9 ± 3.34; range: 41.6-65.2; 39.1-56.8; p > 0.05) (zolpidem + CBD - 52.3 ± 5.03; range: 39.5-73.5; p > 0.05). The mean median decay times are trended towards increase with 1*µ*M zolpidem though neither this showed significance (zolpidem - 8.93 ± 0.54 – 11.0 ± 0.19 – 11.0 ± 0.40; range: 7.00-11.0; 10.6-11.9; 9.80-12.2). A p value of 0.052 overall was returned confirming no significant effect, with the multiple comparison post-hoc also producing a p value of 0.063 between the baseline and zolpidem + 30μM CBD. Intriguingly, 1*µ*M zolpidem evoked a highly significant increase in the mean NICT value (**Biv;** zolpidem - 187 ± 24.1; range: 128-274; p = 0.0017, with the post-hoc p = 0.0045), which occluded the effects of CBD (**Biv;** zolpidem + CBD - 152 ± 20.3; range: 87.6-214).

## Discussion

### CBD increases sIPSC decay time and inhibitory charge transfer in SE but not AMC rats

Overall, CBD had no significant effect on sIPSCs in control rats but significantly increased inhibitory charge transfer in SE rats. The most consistent component of the altered charge transfer was an increase in the decay time constant, which we also observed in human tissue.

### CBD has similar effects on sIPSC in slices of resected human brain from epilepsy patients

The data obtained from resected human brain tissue taken from patients undergoing surgery for refractory epilepsy (nominally pyramidal cells determined by morphological observations and location within the slice) are largely in agreement with the SE rat tissue data. There were no significant differences after CBD application in the mean median amplitude nor in the mean median IEI in human tissue. There was, however, a significant increase in the current decay times and NICT values, similar to the observations in SE rats. It is striking that statistically significant differences were detected despite the limited and infrequent availability numbers of fresh resected human tissue.

We interpret these data to indicate that:

1. The RISE protocol successfully models key aspects the human epilepsy condition in responses to CBD, which are not present in control age-matched control rat.
2. In the RISE model CBD acts on GABA_A_Rs to increase decay times, reminiscent of a benzodiazepine effect and leading to increased inhibitory charge transfer across the membrane.

### CBD interaction with GABA_A_Rs

The BZ inverse agonists flumazenil and β-carboline blocked the effects of CBD on IEI, decay time and NICT, suggestive of direct interaction between CBD and the BZ site of GABA_A_Rs, however, the effects of CBD were additive with low concentrations of the BZ agonist zolpidem, suggesting that CBD acts via a site distinct from the classical BZ-site. Consistent with this, using a range of GABA_A_R subunit combinations expressed in Xenopus oocytes, it has been reported that CBD still exerts BZ-like effects in recombinant receptors that lack the BZ site (Bakas et al, 2017). Taken together, these data suggest that CBD and BZ both enhance GABA_A_R function in similar ways but that they do not act at the same binding site. CBD also displayed an additive effect with pentobarbital, again suggestive of CBD operating allosterically to augment GABA_A_R function. Thus, the increased decay time elicited by both CBD and BZ suggest that CBD is a PAM that binds the GABA_A_R at an as yet unidentified allosteric site to elicit conformational/functional changes.

A notable feature of CBD is the lack of consistency of response. We hypothesise that this could be due the presence of endogenous benzodiazepine-like ligands (endozopines; Iversen, 1977; Costa and Guidotti, 1985) the concentrations of which vary across slices. One of these, diazepam-binding inhibitor (DBI), has been proposed to be able to act as both a NAM (Guidotti et al., (1983); Alfonso et al., 2012) or a PAM (Christian et al., 2013). We therefore speculate that variable levels of DBI in different slices may bind GABA_A_Rs to alter their functional conformations to modify CBD actions, leading to the variation noted across the single cell CBD experiments.

In conclusion, we have shown that CBD is effective in enhancing GABAergic inhibitory signalling in both human and rat epileptic tissue.

